# Fingolimod Regulates BDNF-AS and HOTAIR Expression in Multiple Sclerosis Patients

**DOI:** 10.1101/2021.08.09.455760

**Authors:** Fatemeh Khani-Habibabadi, Mohammad Ali Sahraian, Mohammad Javan, Mehrdad Behmanesh

## Abstract

Brain-derived neurotrophic factor (BDNF), a member of the neurotrophin family, is expressed by neurons and glial cells in the central nervous system (CNS). In the CNS, BDNF is responsible for neuroprotection and neurogenesis. Recent studies showed that the Fingolimod, the first oral medicine for relapsing-remitting multiple sclerosis (RR-MS), induces BDNF expression. Besides, It is well demonstrated that long noncoding RNAs (lncRNAs) have a pivotal role in gene regulation. This study is mainly focused on how Fingolimod treatment plays role in BDNF regulation in coordination with lncRNAs. An in-silico study was performed to predict BDNF-regulatory candidate lncRNAs using online tools. Then, the expression of BDNF-related lncRNAs was analyzed in patients with relapsing-remitting multiple sclerosis (RRMS) at baseline and after three months of Fingolimod treatment. Based on in silico results, two lncRNAs with potential regulatory functions on the BDNF including, Metastasis Associated Lung Adenocarcinoma Transcript 1 (MALAT1) and HOX Transcript Antisense RNA (HOTAIR), and also natural antisense of BDNF were selected. Fingolimod treatment increased the expression of HOTAIR lncRNA; however, the BDNF antisense RNA (BDNF-AS) expression was reduced dramatically. Furthermore, the results indicate a positive correlation between HOTAIR and MALAT1 lncRNAs and BDNF. Also, after Fingolimod treatment, the patients’ EDSS scores were declined or remained unchanged, indicating disease hindrance by Fingolimod therapy. Altogether, fingolimod exerts protective roles in RRMS patients probably by the mediation of HOTIAR and BDNF-AS lncRNAs.

## 1. Introduction

Multiple Sclerosis (MS) is an inflammatory autoimmune disease in which myelin sheets wrapping central nervous system (CNS) neurons are exposed to degradation due to the inflammatory milieu resulting in patients’ cognitive and motor impairments. In the CNS, myelin loss makes the naked axons vulnerable to pro-inflammatory cytokines (Thompson et al. 2018). Based on clinical symptoms, MS divides into four subgroups, clinically isolated syndrome (CIS), relapsing-remitting MS (RRMS), primary progressive MS (PPMS), and secondary progressive MS (SPMS). In most MS patients, the disease occurs as the relapsing-remitting phase that usually proceeds to the chronic progressive stage (Barnett and Prineas 2004).

The inflammatory process in MS pathogenesis is dependent on lymphocytes migration out of the lymph nodes and crossing through the blood-brain barrier (BBB) into the CNS. A G protein-coupled receptor, sphingosine 1-phosphate receptor 1 (S1P1), is involved in the migration of lymphocytes to the circulation based on the S1P gradient. Sphingosine activates S1P and triggers several ubiquitous pathways through the body including, Mitogen-Activated Protein Kinase (MAPK), cAMP response element-binding (CREB), Phospholipase C (PLC), Protein kinase C (PKC), Cyclic adenosine monophosphate (cAMP), and calcium signaling pathway (Dev et al. 2008).

The S1Ps can be targeted by sphingosine analog, Fingolimod (FTY-720). Like sphingosine, Fingolimod must become phosphorylated by sphingosine kinase (SphK) to convert to its active form (FTY720-P) (Geffin et al. 2017). Phosphorylated Fingolimod can bind to all S1P subtypes (S1P1, S1P3, S1P4, and S1P5) except for S1P2. The binding of Fingolimod to the S1P1 receptor induces its internalization and endosomal degradation, thereby inhibiting its response to the S1P gradient and lymphocyte egress out of the lymph nodes. This mechanism is called the functional antagonist (Pelletier and Hafler 2012). On the other hand, an increasing body of evidence suggests that fingolimod plays roles in remyelination and neuroprotection, by crossing through the BBB and binding to S1P receptors on microglia, oligodendrocytes, astrocytes, and neurons cells (Deogracias et al. 2012) (Pelletier and Hafler 2012; Kappos et al. 2006). Furthermore, it is declared that fingolimod induces brain-derived neuroprotective factor (BDNF) expression through MAPK and CREB phosphorylation (Deogracias et al. 2012).

BDNF is a member of the neurotrophin family, with proved endogenous neuroprotective function which also plays roles in myelin repair in the demyelinating diseases’ models (Fulmer et al. 2014). BDNF involves in CNS development through the control of neuronal survival, differentiation, and protection (Almeida et al. 2005). BDNF is secreted not only by neurons but also by different cell type, e.g., in the active and inactive multiple sclerosis lesions, infiltrated B cell, T cells, and also macrophages play a neuroprotective role by producing BDNF (Kerschensteiner et al. 1999; Stadelmann et al. 2002). More than its protective effects on neurons, BDNF induces oligodendrocytes proliferation, differentiation, and remyelination (Fletcher et al. 2018; Xiao et al. 2013).

Long non-coding RNAs (lncRNAs) are a category of non-coding RNAs with more than 200 nucleotides length that involve in either nuclear or cytoplasmic compartments. Through miRNAs sponge, protein decoy, and intracellular scaffolds, lncRNAs could play their role in gene regulation at different stages including, transcription, post-transcription, and protein translation. Recent studies have demonstrated the potential roles of lncRNAs in neurogenesis, rapid CNS development, neural functions, and development of neurodegenerative disorders (Yang et al. 2018; Wu et al. 2013).

Considering the pivotal role of BDNF in neuroprotection, understanding its regulatory mechanisms could be essential in resolving the neurobiology and management of neural disorders. Here, considering the ambiguous functions of lncRNAs in the BDNF regulation, the regulatory roles of HOX Transcript Antisense RNA (HOTAIR), Metastasis Associated Lung Adenocarcinoma Transcript 1 (MALAT1), and BDNF antisense RNA (BDNF-AS) lncRNAs on BDNF expression were studied. By in silico study and literature review, these three lncRNAs were selected, and then their expression was quantified in RRMS patients at baseline and after three months of Fingolimod treatment. Also, the EDSS score and its correlation with these gene expression levels were measured. This is the first study that evaluates the Fingolimod effects on the lncRNAs expression concerning BDNF expression in RRMS patients.

## 2. Materials and methods

### 2.1. Selecting candidate lncRNAs

To candidate BDNF-regulating lncRNAs, we employed online software starBase v2.0 (J.-H. Li et al. 2014) to shown RNA binding proteins and miRNAs related to genes transcripts, ChiPBase v2.0 (K.-R. Zhou et al. 2016) and FARNA (Alam et al. 2017) to obtain transcription factors binding sites, lncRNA2Target to demonstrate lncRNAs-related genes by knockdown experiments. To understand whether lncRNAs could be differentially regulated under Fingolimod treatment, the promoter regions of *BDNF* and candidate lncRNAs were scanned by the JASPAR database to discover binding sites of these inducible transcription factors.

The promoter regions of *BDNF* and candidate lncRNAs of MALAT and HOTAIR were scanned by JASPAR database to discover binding sites of inducible transcription factors of nuclear factor of activated T-cells 1 (NFAT1), activator protein 1 (AP1), nuclear factor kappa B (NFκB) and CREB activation (Baer et al. 2018; Deogracias et al. 2012).

### 2.2 Patients and sample collection

Participants were selected from 20 female relapsing-remitting MS patients referred to the MS Research Center of Sina Hospital of Tehran University of Medical Sciences, Tehran, Iran, from February 2018 to May 2019. All of the patients were clinically diagnosed according to McDonald’s criteria by an expert neurologist.

The time of blood sampling (2 ml) was at the baseline (within two weeks after their previous relapse) and three months after Fingolimod therapy (Danelvin®). After three months of Fingolimod therapy, ten patients admitted resuming their participation in the research. The exclusion criteria defined as pregnancy, lactation, and smoking. Three patients were excluded from the study, due to their detrimental side effects to the Fingolimod therapy. Ten healthy individuals also were participated as the control group (Table 1). All of the participants signed the consent letter, and the steps were done based on the Helsinki declaration in research on human samples. The study was approved by the ethical committee of Tarbiat Modares University (ID: IR.TMU.REC.1396.607).

**Table 1.**
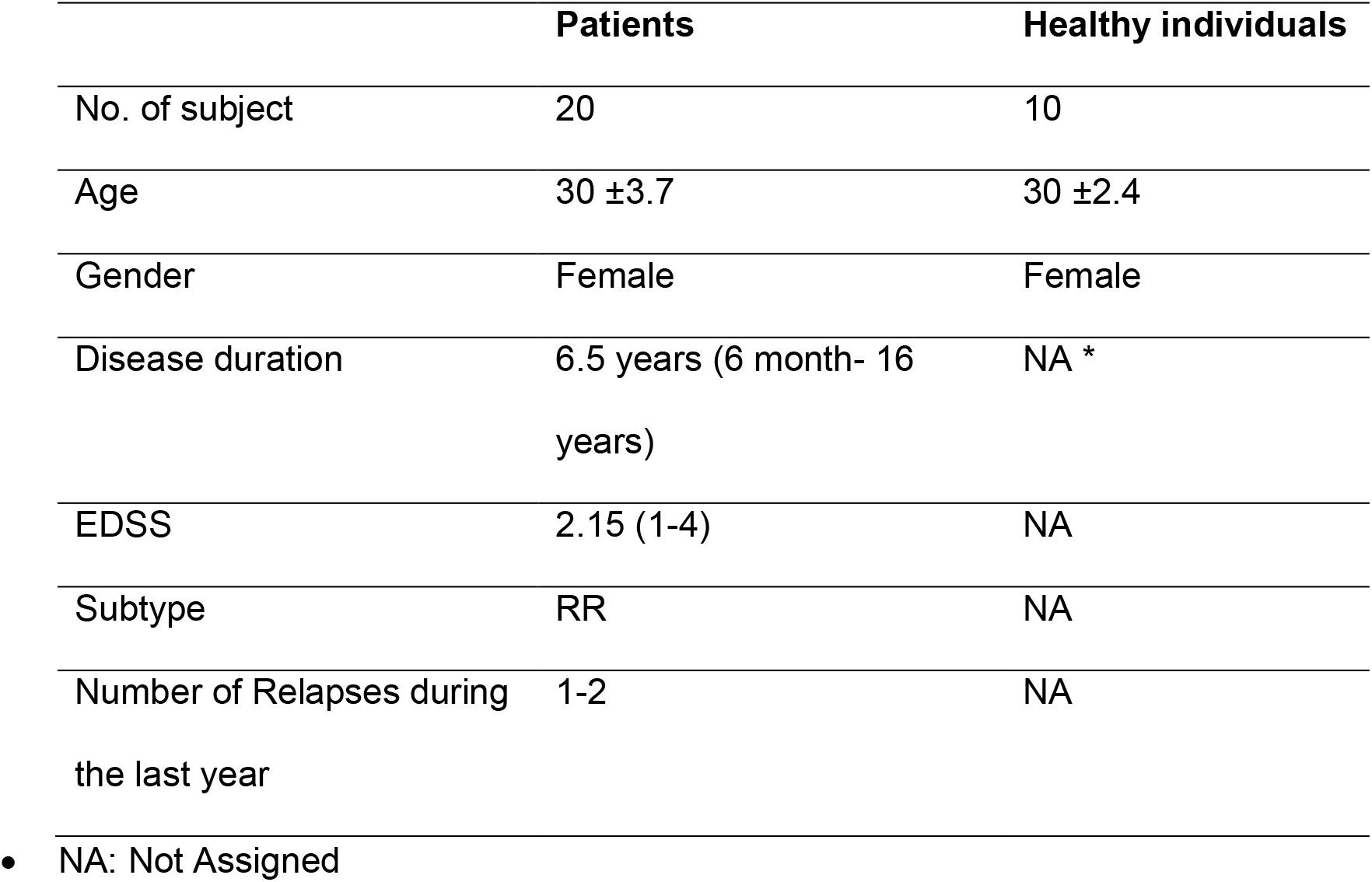
Demographic Features of MS patients and control group. Twenty women with MS who referred to the MS research Center of Sina Hospital for receiving the first dose of Fingolimod enrolled in the study. Ten age, sex and ethnicity healthy individuals were selected as the control group.

### 2.3 RNA extraction and gene expression analysis

PBMCs were isolated from fresh blood samples by Ficoll gradient centrifugation technique (lympholyte, Cedarlane, Netherlands). PBMCs total RNA was extracted using the RiboEX solution (GeneAll, Seoul, Korea), and its quality and quantity were measured by spectrophotometry and 1.5% agarose gel. DNaseI (Fermentas, USA) treatment was performed on RNA samples to remove genomic DNA contamination. Three µg of total RNAs was applied to synthesize the first strand of cDNA using M-MulV reverse transcriptase (Thermo Scientific, USA), oligo dT and random hexamer primers (MWG, Germany) in a total of 20 µl reaction mixture according to manufacturer’s protocol.

### 2.4 Primer design and gene expression analysis

For gene expression analysis, specific primers were designed by NCBI primer blast and Oligo 7 software. The sequences of designed primers are demonstrated in table 2. Quantitative real-time PCR was performed using a StepOne™ sequence detection system (Applied Biosystems, USA), with 10 ng cDNA, 5 µl of 2X SYBR® Green master mix (Solis BioDyne, Estonia) and 200 nm of each forward and reverse primers up to final reaction volumes of ten µl. Differential gene expression analysis was performed using the 2^-ΔCT^ method (Livak and Schmittgen 2001). The expression level of target genes was normalized to the expression of GAPDH as the internal control, and each sample was assayed at least in duplicate.

**Table 2.**
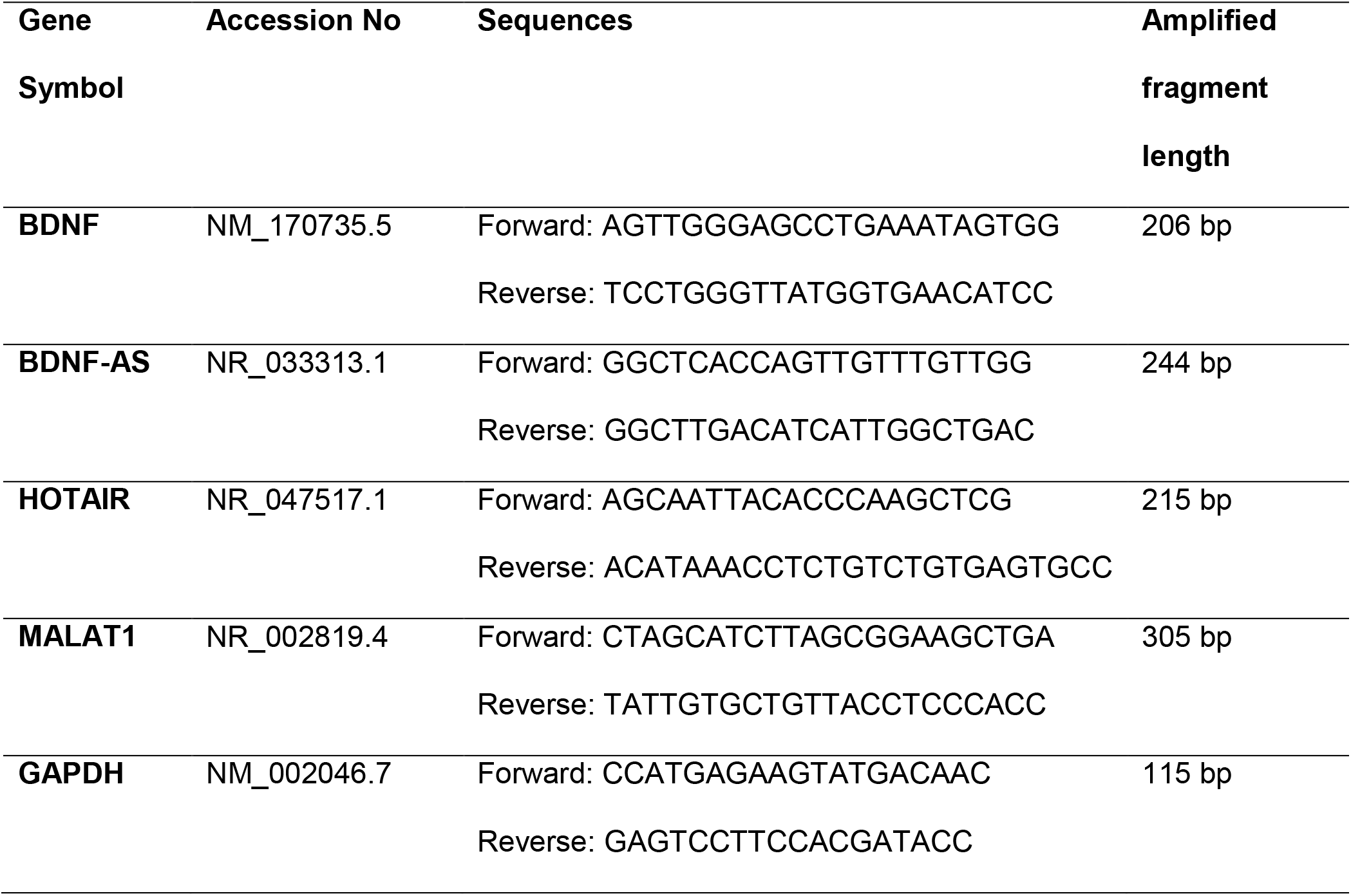
Primer pairs’ sequences used for analyzing gene expression by RT-qPCR.

### 2.5 Statistical analysis

For multiple-groups analysis in non-Gaussian distributions, Kruskal-Wallis with Dunn’s correction was performed. A comparison between paired groups was done using Wilcoxon matched pairs. The Spearman test was performed to explore correlated genes. All of the statistical analyses and graphs visualizations were accomplished by Prism – GraphPad 7 (GraphPad Software, USA) and R-3.5.1 software. Error bars represent median with significance level was assigned as p < 0.05 (two-tailed test).

## 3. Results

### 3.1. Selection of three candidate lncRNAs with potential roles in regulating BDNF gene expression

To predict candidate BDNF-regulatory lncRNAs, sponging BDNF-regulatory miRNAs, interaction with chromatin modulating complexes, expression, and recruitment of transcription factors to the BDNF promoter region were employed. Table 3 represents candidate lncRNAs play roles in BDNF expression level regulation through various mechanisms. MALAT1 and HOTAIR lncRNAs were selected as candidate lncRNAs for the regulation of the BDNF expression level using online software. Furthermore, BDNF-AS was included in the study because of its validated function on BDNF promoter methylation and initiation of transcription (Modarresi et al. 2012; Bohnsack et al. 2019).

**Table 3.**
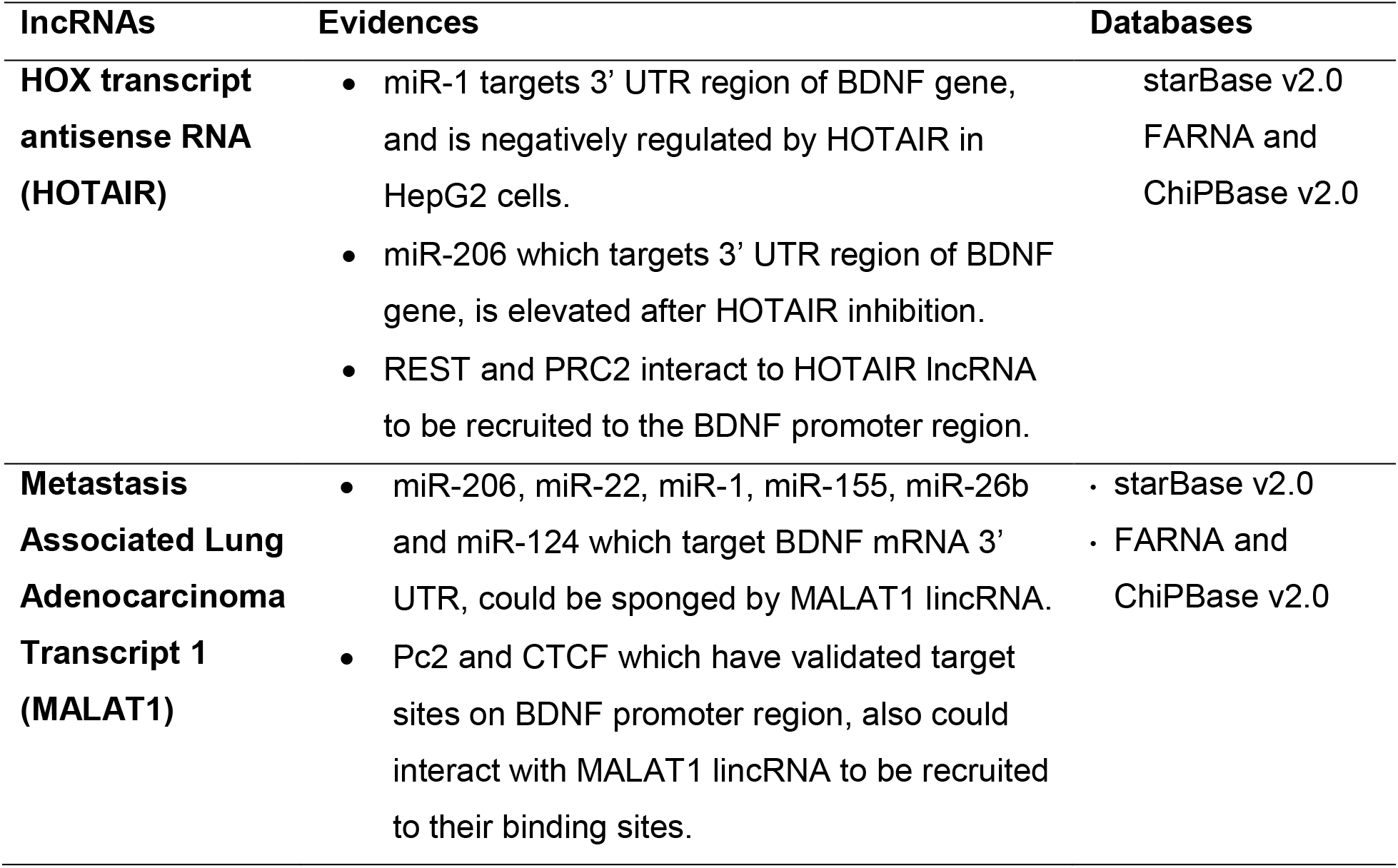
Predicted lncRNAs with potential regulatory function on BDNF expression.

### 3.2. BDNF, BDNF-AS, and candidate lncRNAs probably are induced by Fingolimod treatment

To investigate whether the candidate lncRNAs can be induced by fingolimod treatment or not, promoter region (5 Kb upstream and 1 Kb downstream) of these lncRNAs were scanned to find the binding sites for transcription factors that are confirmed to be activated by fingolimod treatment including, NFAT1, NfkB, AP1, and CREB (Deogracias et al. 2012; Baer et al. 2018). For algorithm similarity, the minimum relative profile score threshold assigned 85%. All of these factors had at least one binding site on BDNF, BDNF-AS, and the candidate lncRNAs promoter region, which demonstrates their possible regulation by fingolimod treatment (Table 4). Taken together, in silico analysis showed probable binding sites of NFkB, NFAT1, CREB, and AP1 transcription factors on scanned promoters. Hence, these genes probably can be induced by Fingolimod treatment.

**Table 4.**
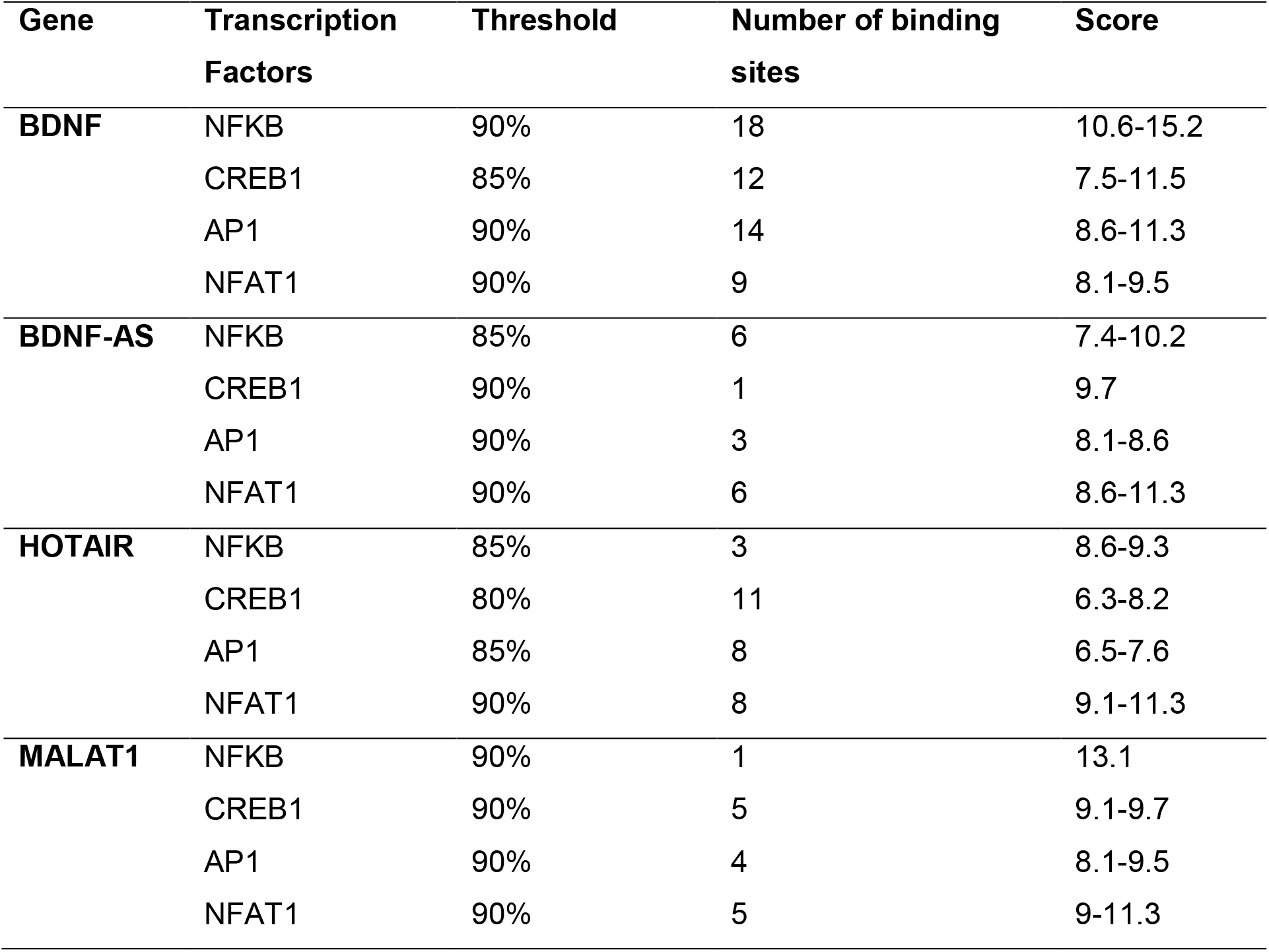
The results of scanning the promoter regions of BDNF and selected lncRNAs for possible binding sites of NFKB, CREB1, AP1, and NFAT1.

| **Gene** | **Transcription Factors** | **Threshold** | **Number of binding sites** | **Score** |
| --- | --- | --- | --- | --- |
| **BDNF** | NFkB | 90% | 18 | 10.6-15.2 |
|  | CREB1 | 85% | 12 | 7.5-11.5 |
|  | AP1 | 90% | 14 | 8.6-11.3 |
|  | NFAT1 | 90% | 9 | 8.1-9.5 |
| **BDNF-AS** | NFkB | 85% | 6 | 7.4-10.2 |
|  | CREB1 | 90% | 1 | 9.7 |
|  | AP1 | 90% | 3 | 8.1-8.6 |
|  | NFAT1 | 90% | 6 | 8.6-11.3 |
| **HOTAIR** | NFkB | 85% | 3 | 8.6-9.3 |
|  | CREB1 | 80% | 11 | 6.3-8.2 |
|  | AP1 | 85% | 8 | 6.5-7.6 |
|  | NFAT1 | 90% | 8 | 9.1-11.3 |
| **MALAT1** | NFkB | 90% | 1 | 13.1 |
|  | CREB1 | 90% | 5 | 9.1-9.7 |
|  | AP1 | 90% | 4 | 8.1-9.5 |
|  | NFAT1 | 90% | 5 | 9-11.3 |

### 3.3. BDNF expression is elevated in MS patients

To check the expression level of studied genes in MS patients, the expression of BDNF, BDNF-AS, and the candidate lncRNAs was analyzed in control and 20 RRMS patients at initiation of fingolimod therapy (baseline), . Gene expression study demonstrated elevated BDNF expression levels in RRMS patients comparing the control group (fold= 4.2, p=0.027). Given the BDNF expression level, patients were categorized into two groups, a group with similar BDNF expression to the healthy individuals (9 patients) and the other group with higher BDNF expression (11 patients). The demographic features of MS patients revealed that the mean of the EDSS score in the group with common BDNF expression was 1.5, but it was 2.8 in the other group. Also, the disease duration was 5.3 years in the group with common BDNF expression; however, it was 7.8 years in the group with a higher BDNF expression level (Figure 1, A). In contrast, the BDNF-AS expression level did not differ between the control group and MS patients (fold= -1.2, p=0.939) (Figure 2, B). Also, gene expression analysis revealed no alteration in HOTAIR lncRNA expression level between MS patients and healthy individuals (fold= 2.2, p=0.108) (Figure 1, C). Similarly, the MALAT1 expression level in MS patients was not significantly upregulated compare to the control group (fold= -2.4, p=0.074), though the wide variation in expression level in the patients (Figure 1, D).

**Fig. 1.**
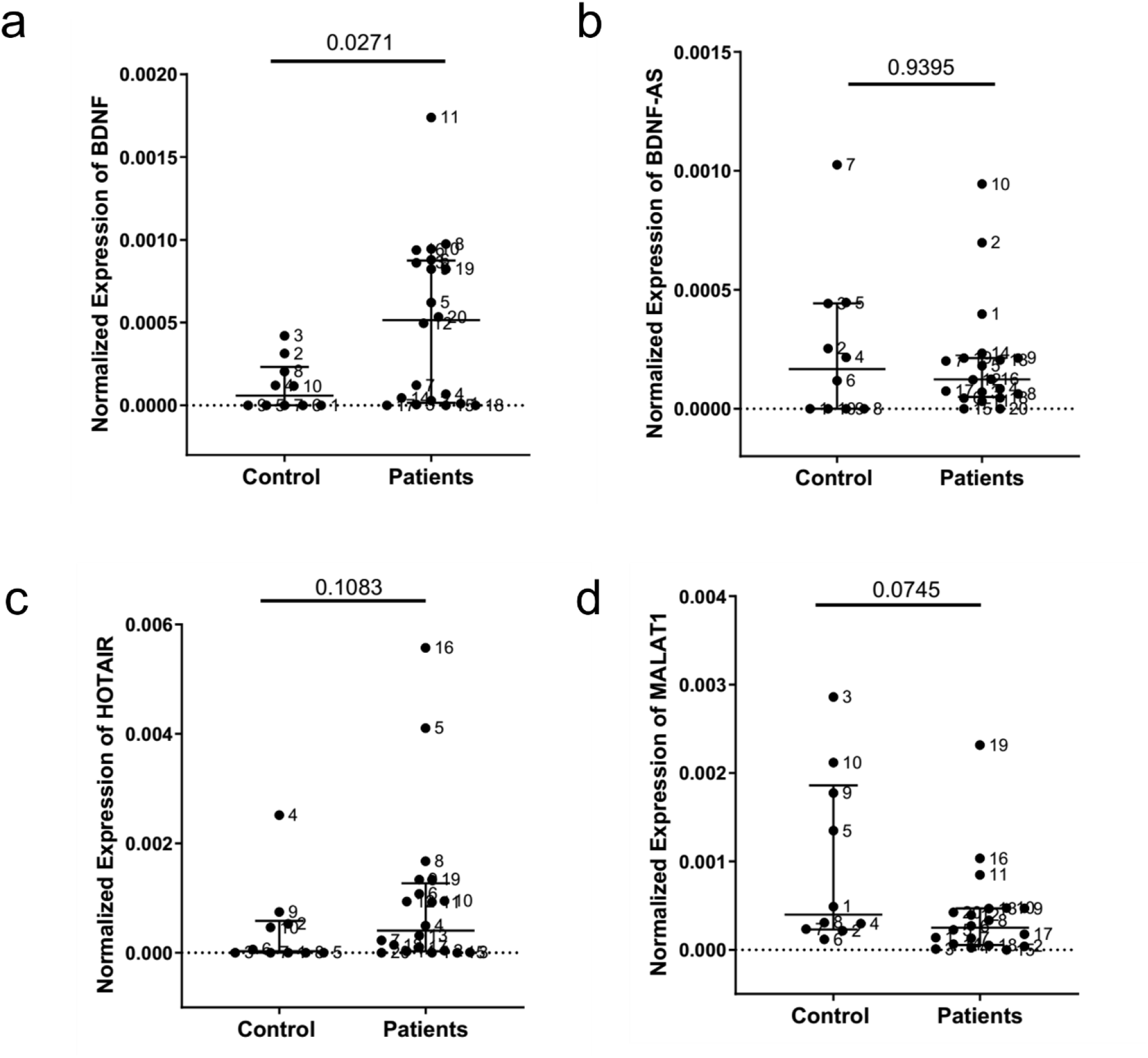
Evaluation of gene expression level in RRMS patients at baseline. (A) The expression level of BDNF in MS patients (n = 20) is higher than the control group (P = 0.027). (B) The level of BDNF antisense expression was not significantly different between the control and patient groups (P = 0.939). (C) The expression level of HOTAIR lncRNA was not significantly different between the patient and healthy groups (P = 0.108). (D) Similarly, the expression level of MALAT1 lncRNA in MS patients did not show a significant decrease compared to the control group (P = 0.074). The significance threshold was considered p <0.05. Error bar = Median with interquartile range.

**Fig. 2.**
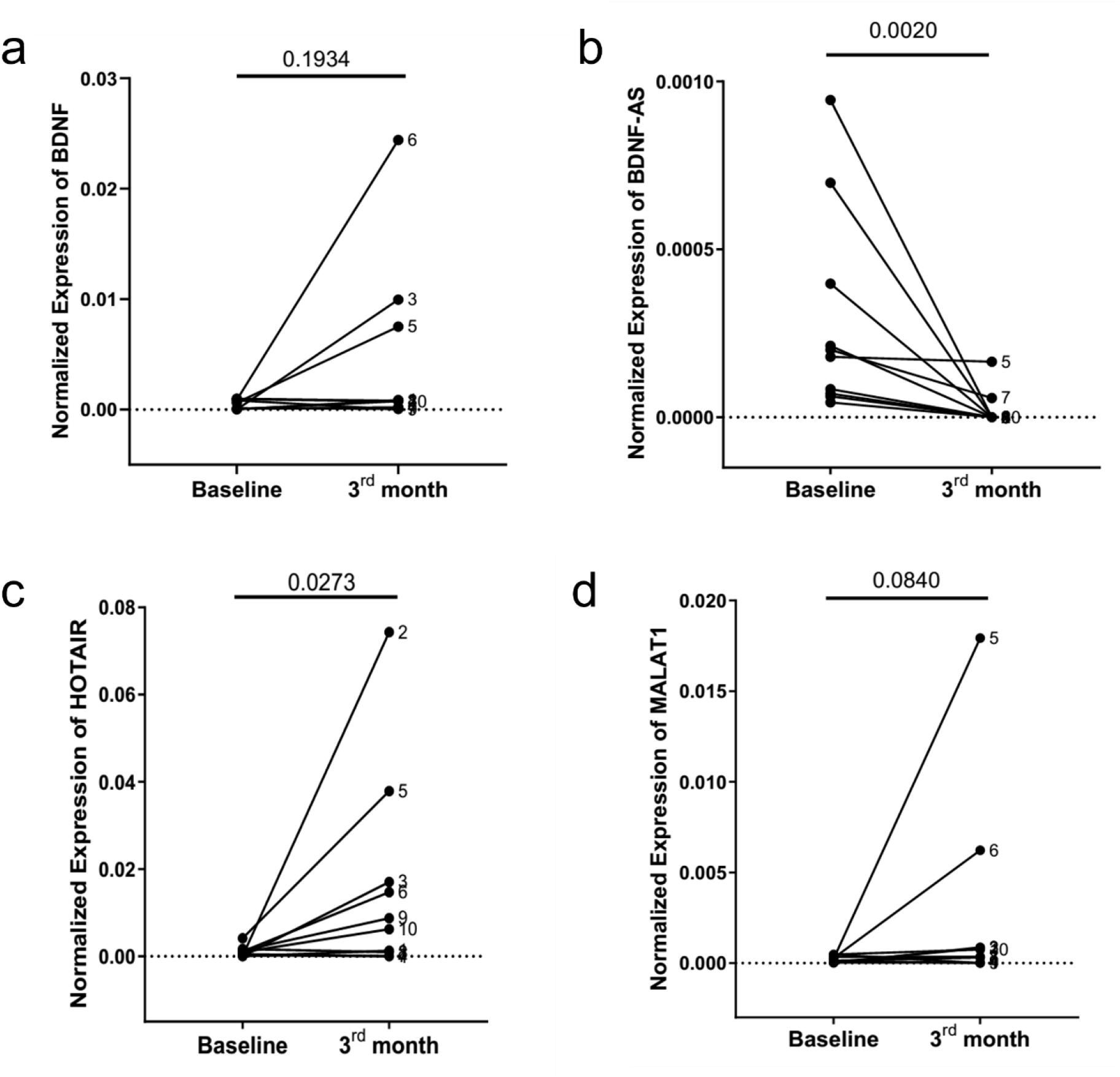
The effects of fingolimod treatment on the expression level of the studied genes. (a) three months of fingolimod treatment did not affect BDNF gene expression level (P = 0.83); (b) However, After three months of taking medication, the expression level of BDNF-AS declined sharply (P = 0.002). (c) But, fingolimod therapy significantly increased the expression level of HOTAIR lncRNA (P = 0.027). (d) Similar to the BDNF, treatment with fingolimod did not affect MALAT1 gene expression (p = 0.084). The significance threshold was considered p <0.05. Error bar = Median with interquartile range.

### 3.4. Fingolimod significantly reduced BDNF antisense level

Patients’ PBMCs gene expression analyses were redone in the same patients after three months of fingolimod therapy to monitor the medicine effects on the candidate genes expression level. 10 patients were excluded from the study because they did not wish to continue their participation or clinical manifestation of severe side effects. As shown in Figure 2, A, three months of treatment with Fingolimod did not significantly alter the expression level of the gene, despite a sharp increase in BDNF expression in three patients (fold = 5.57, p = 0.193). The distribution of BDNF gene expression after three months of drug administration differs notably from the median (Figure 2, A). In contrast, BDNF antisense expression levels decreased greatly after three months of Fingolimod administration (fold = -12.9, p = 0.002) (Figure 2, B). In contrast, the expression level of lncRNA HOTAIR increased significantly with three months of Fingolimod treatment (fold = 10.49, p = 0.027) (Figure 2, C). However, Fingolimod administration did not significantly alter the expression level of MALAT1 lncRNA (fold = 3.2, p = 0.084) (Figure 2, D).

### 3.5. Fingolimod altered the gene correlations

Next, correlation analyses were performed between these groups to investigate their possible interactions and also the fingolimod effects on these relationships. In healthy individuals, there was no correlation between BDNF and candidate lncRNAs; however, in MS patients at baseline, there was a significant correlation (n = 10) between BDNF gene and HOTAIR (r = 0.80, p = 0.007) and MALAT1 lncRNAs (r = 0.85, p = 0.002) (Fig. 3, a & b). After three months of fingolimod administration, the correlation between lncRNAs and the BDNF was lost, and a new relationship was obtained between these two lncRNAs (r = 0.67, p = 0.037) (Fig. 3, c). These results indicate the effect of Fingolimod medicine on the correlation of these genes.

**Fig. 3.**
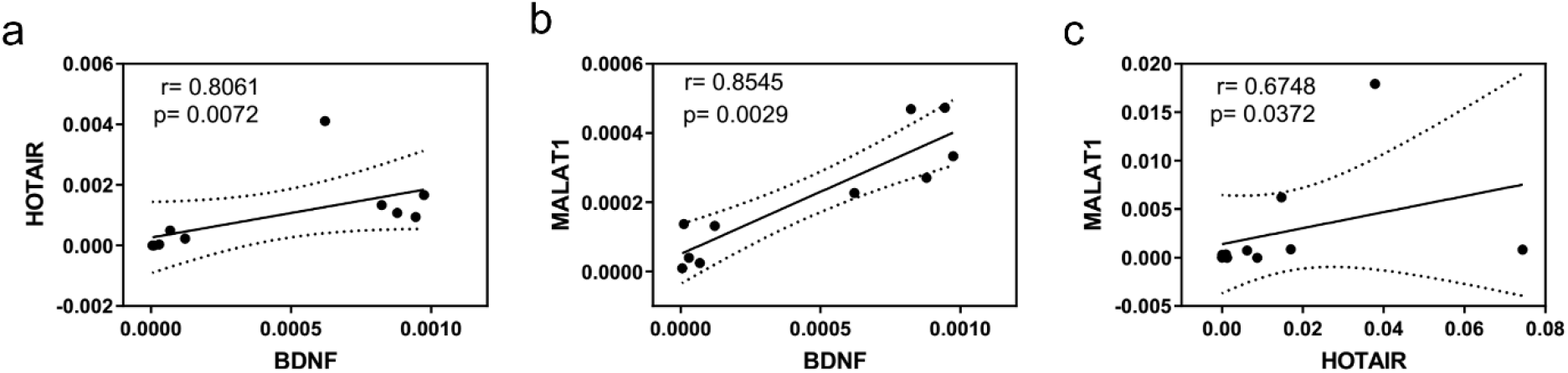
Investigating the correlation between gene expression levels at baseline and three months of therapy to find possible intergenic connections. (a) In MS patients (n = 10) before the start of treatment, a significant positive correlation was observed between BDNF gene expression and HOTAIR (r = 0.80). (b) Also, a significant positive correlation was observed between BDNF gene expression and MALAT1 (r = 0.85). (c) Three months of Fingolimod administration, while eliminating the association of the BDNF gene with HOTAIR and MALAT1, resulted in a significant positive correlation between these two lncRNAs. The significance threshold was considered p <0.05.

### 3.6 Fingolimod treatment hindered disease progression

The EDSS score is an indicator of the disease progression in MS patients. Here, to measure the fingolimod effects on disease progression, the EDSS score was checked at baseline and one year after fingolimod administration. The results showed that the severity of the disability remained steady in four patients, while in the other patients, the EDSS score decreased about 0.5 to 1 unit (Fig. 4, a). The statistical study showed a significant decrease in the EDSS score after fingolimod administration (p = 0.031) (Fig.4, b). Correlation analyses revealed no relationship among the EDSS score, BDNF gene, and candidate lncRNAs. The rate of EDSS score at baseline and the duration of the disease varied between patients who had a decrease in EDSS score (group A) and those who did not show a reduction in the EDSS score after fingolimod treatment (group B). The initial EDSS score in group A was 2.25, but in group B, this score was 1.25 (Fig. 4, c). Also, the duration of the disease in group A was 7.9 years, and in group B, it was 4.5 years (Fig. 4, d). These results indicate the importance of EDSS score and disease duration on the fingolimod effects on clinical manifestation.

**Fig. 4.**
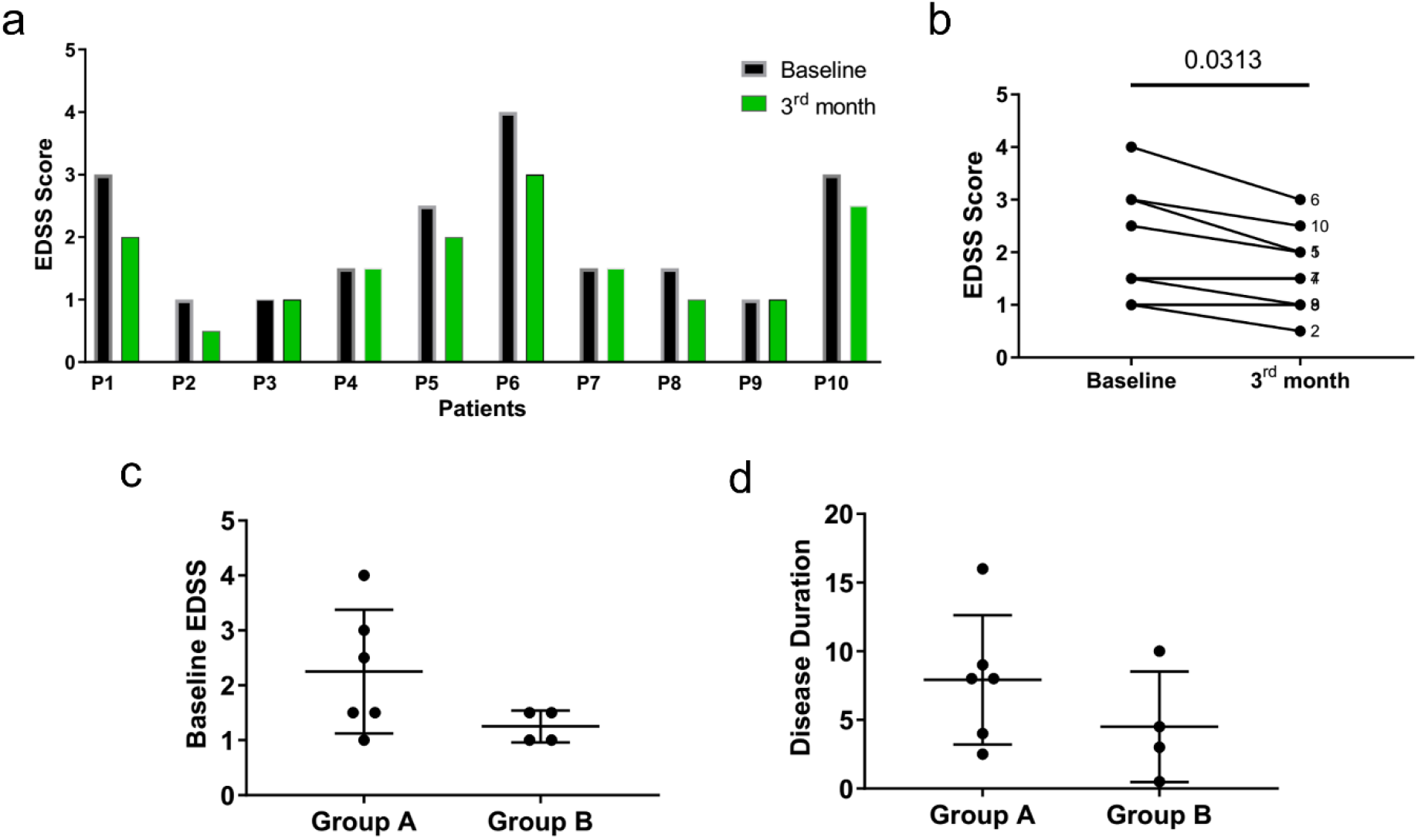
The effects of the fingolimod treatment on the EDSS score. (a) In MS patients (n = 10), the fingolimod reduced the EDSS score in 60% of patients (group A), and the rest 40% of patients (group B) hindered the progression of the disease. (b) Analysis of the EDSS score before and after taking fingolimod revealed a significant decrease in the EDSS score after the treatment. (c) Though in group A the mean baseline EDSS score was 2.25, this score was 1.25 in the B group. (d) The mean disease duration in group A was 7.9 years, while the disease duration in group B was 4.5 years. The significance threshold was considered p <0.05. Error bar = Median with interquartile range.

## 4. Discussion

According to the BDNF fundamental roles in neurons’ survival and myelination, it seems that deciphering BDNF regulatory mechanisms can pave the way toward new approaches for novel therapeutic strategies. Given the pivotal roles of lncRNAs in gene regulation of neurodegenerative disease (Wu et al. 2013), the function of these noncoding molecules in the BDNF regulation was investigated in this study. Here, the BDNF expression was higher in MS patients who had referred to the MS Center of Sina Hospital to receive fingolimod within two weeks after experiencing their previous relapse. Similar to this result, during the relapse phase of RRMS patients or within two weeks after it, the serum level of BDNF significantly elevated, comparing the remission phase. The overexpressed BDNF by infiltrated lymphocytes, neurons, and glial cells causes oligodendrocyte precursor cell proliferation, remyelination of plaque, and axonal regeneration (Frota et al. 2009; Caggiula et al. 2005; McTigue et al. 1998). Given the vital role of BDNF in the brain, increased BDNF expression in these patients may be involved in the repair process. Furthermore, patients who had higher expression of BDNF than the control also had higher EDSS scores and disease duration comparing other patients. These results suggest that the more advanced the disease, the higher the amount of BDNF produced in the lymphocyte cell, possibly to compensate or repair the damage according to its neuroprotective role.

The results showed a significant decrease in BDNF-AS expression after three months of fingolimod administration. Previously, it has been reported that the BDNF-AS lncRNA suppresses the BDNF promoter region by H3K27me3 through the recruitment of polycomb repressive complex 2 (PRC2) and enhancer of zeste homolog 2 (EZH2), also its Knockdown increased BDNF expression (Modarresi et al. 2012; Bohnsack et al. 2019). Base on the BDNF-AS suppressing role on BDNF expression, the downregulation of BDNF-AS by fingolimod treatment may play a role in the positive regulation of BDNF. Here, the lack of BNDF overexpression by fingolimod may be a consequence of the limited number of participants. Also, it was declared that BDNF-AS could play a pro-inflammatory role by inducing TNF-α and also suppressing IL-2 anti-inflammatory pathways (Xu et al. 2016). On the other hand, fingolimod indirectly reduces the production of inflammatory cytokines, including IL-6, TNF-α, IL-1β hence has an anti-inflammatory function (Lee et al. 2017; Bohnsack et al. 2019). Given the BDNF-AS inflammatory roles and the anti-inflammatory function of fingolimod, this medicine may exert its anti-inflammatory function by suppressing the expression of BDNF-AS.

This study revealed that fingolimod increased HOTAIR expression simultaneously with the hindrance of disease progression. In support of this finding, previous studies suggest protective roles for HOTAIR in inflammatory diseases including CAD, Diabetes, and rheumatoid arthritis through mediating NF-κB pathway, oxidative damage, inflammation, and cell death (Xu et al. 2016). Altogether, this data may suggest protective roles for Hotair in MS patients in response to fingolimod treatment.

Furthermore, here it was demonstrated that BDNF is correlated with MALAT1 lncRNA and possibly linked to each other by intracellular signaling pathways. It has been shown that binding of the CREB transcription factor to MALAT1 lncRNA stabilizes its phosphorylated form and inducing its binding to the BDNF gene promoter region (Yao et al. 2016) (Deogracias et al. 2012). Hence MALAT1 is probably associated with the regulation of BDNF through CREB stabilization.

Another interpretation for the positive correlation among BDNF, HOTAIR, and MALAT1 lncRNAs is the competitive endogenous RNA (ceRNA) hypothesis. The hypothesis of ceRNA is based on the interaction among lncRNAs, mRNAs, and miRNAs through microRNA binding sites (MREs) on lncRNAs. According to this hypothesis, mRNA and lncRNA could influence the expression of each other through competition for binding to the restricted miRNA pool (Salmena et al. 2011). Previous studies have provided data that can support the possible interaction of HOTAIR and MALAT1 with BDNF regulation via the ceRNA mechanism. Also, the ceRNA hypothesis explains why a significant positive correlation between MALAT1 and HOTAIR lncRNAs may exist (Tian et al. 2014; Khani-Habibabadi et al. 2019, Q. Li et al. 2018; Chang et al. 2018) However, the cause of this positive correlation between HOTAIR and MALAT1 needs further study.

This study showed that a year of fingolimod treatment had a positive effect in ameliorating disabilities and symptoms by a reduction in or the maintenance of EDSS score and inhibiting a recurrence of relapse. Similarly, previously it has been represented that taking Fingolimod decreases EDSS score and the number of relapses in most patients (Mazibrada et al. 2018; Mazdeh et al. 2019). These data suggest protective roles for fingolimod in MS alleviation. Also, it seems that patients’ response to fingolimod treatment is related to the initial EDSS score, disease duration, and the genetic background of individuals. So it is necessary to pay attention to demographic characteristics and personalized medicine in using this drug in the treatment of MS patients.

## 5. Conclusion

In this study, HOTAIR and MALAT1 regulatory roles on BDNF expression and the effect of fingolimod on their interaction were assessed. The results showed that after a relapse in MS patients, the level of BDNF expression is higher than in healthy individuals. Also, it showed that fingolimod decreased BDNF-AS and increased HOTAIR expression simultaneously with disease hindrance probably resulted in a less inflammatory response. Furthermore, the results indicate a positive regulation of HOTAIR and MALAT1 on BDNF expression, and also between HOTAIR and MALAT1 lncRNAs. These connections are probably caused by the ceRNA and miRsponge mechanisms. Finally, it was suggested that patients’ response to Fingolimod treatment is related to the EDSS score, disease duration, and patients’ genetic backgrounds. To summarize, fingolimod may exert its protective roles in RRMS patients by the regulation of HOTIAR and BDNF-AS lncRNAs.

## Acknowledgements

The authors gratefully appreciate the contributions of the MS patients and healthy controls for blood donations.

## Ethics approval

The study was approved by the ethical committee of Tarbiat Modares University (ID: IR.TMU.REC.1396.607).

## References

Alam, T., Uludag, M., Essack, M., Salhi, A., Ashoor, H., Hanks, J. B., et al. (2017). FARNA: knowledgebase of inferred functions of non-coding RNA transcripts. Nucleic acids research, 45(5), 2838–2848.

Almeida, R., Manadas, B., Melo, C., Gomes, J., Mendes, C., Graos, M., et al. (2005). Neuroprotection by BDNF against glutamate-induced apoptotic cell death is mediated by ERK and PI3-kinase pathways. Cell death and differentiation, 12(10), 1329.

Baer, A., Colon-Moran, W., & Bhattarai, N. (2018). Characterization of the effects of immunomodulatory drug fingolimod (FTY720) on human T cell receptor signaling pathways. Scientific reports, 8(1), 10910.

Barnett, M. H., & Prineas, J. W. (2004). Relapsing and remitting multiple sclerosis: pathology of the newly forming lesion. Annals of neurology, 55(4), 458–468.

Bhan, A., Hussain, I., Ansari, K. I., Kasiri, S., Bashyal, A., & Mandal, S. S. (2013). Antisense transcript long noncoding RNA (lncRNA) HOTAIR is transcriptionally induced by estradiol. Journal of molecular biology, 425(19), 3707–3722.

Bohnsack, J. P., Teppen, T., Kyzar, E. J., Dzitoyeva, S., & Pandey, S. C. (2019). The lncRNA BDNF-AS is an epigenetic regulator in the human amygdala in early onset alcohol use disorders. Translational psychiatry, 9(1), 34.

Brunkhorst, R., Vutukuri, R., & Pfeilschifter, W. (2014). Fingolimod for the treatment of neurological diseases—state of play and future perspectives. Frontiers in cellular neuroscience, 8, 283.

Caggiula, M., Batocchi, A., Frisullo, G., Angelucci, F., Patanella, A., Sancricca, C., et al. (2005). Neurotrophic factors and clinical recovery in relapsing-remitting multiple sclerosis. Scandinavian journal of immunology, 62(2), 176–182.

Cao, S., Wang, Y., Li, J., Lv, M., Niu, H., & Tian, Y. (2016). Tumor-suppressive function of long noncoding RNA MALAT1 in glioma cells by suppressing miR-155 expression and activating FBXW7 function. American journal of cancer research, 6(11), 2561.

Caputo, V., Sinibaldi, L., Fiorentino, A., Parisi, C., Catalanotto, C., Pasini, A., et al. (2011). Brain derived neurotrophic factor (BDNF) expression is regulated by microRNAs miR-26a and miR-26b allele-specific binding. PloS one, 6(12), e28656.

Chandrasekar, V., & Dreyer, J.-L. (2009). microRNAs miR-124, let-7d and miR-181a regulate cocaine-induced plasticity. Molecular and Cellular Neuroscience, 42(4), 350–362.

Chang, L., Guo, R., Yuan, Z., Shi, H., & Zhang, D. (2018). LncRNA HOTAIR regulates CCND1 and CCND2 expression by sponging miR-206 in ovarian cancer. Cellular Physiology and Biochemistry, 49(4), 1289–1303.

Chen, Y.-J., Wang, Y.-N., & Chang, W.-C. (2007). ERK2-mediated C-terminal serine phosphorylation of p300 is vital to the regulation of epidermal growth factor-induced keratin 16 gene expression. Journal of Biological Chemistry, 282(37), 27215–27228.

Deogracias, R., Yazdani, M., Dekkers, M. P., Guy, J., Ionescu, M. C. S., Vogt, K. E., et al. (2012). Fingolimod, a sphingosine-1 phosphate receptor modulator, increases BDNF levels and improves symptoms of a mouse model of Rett syndrome. Proceedings of the National Academy of Sciences, 109(35), 14230–14235.

Dev, K. K., Mullershausen, F., Mattes, H., Kuhn, R. R., Bilbe, G., Hoyer, D., et al. (2008). Brain sphingosine-1-phosphate receptors: Implication for FTY720 in the treatment of multiple sclerosis. Pharmacology & Therapeutics, 117(1), 77–93, doi:10.1016/j.pharmthera.2007.08.005.

Fletcher, J. L., Wood, R. J., Nguyen, J., Norman, E. M., Jun, C. M., Prawdiuk, A. R., et al. (2018). Targeting TrkB with a brain-derived neurotrophic factor mimetic promotes myelin repair in the brain. Journal of Neuroscience, 38(32), 7088–7099.

Frota, E. R. C., Rodrigues, D. H., Donadi, E. A., Brum, D. G., Maciel, D. R. K., & Teixeira, A. L. (2009). Increased plasma levels of brain derived neurotrophic factor (BDNF) after multiple sclerosis relapse. Neuroscience letters, 460(2), 130–132.

Fulmer, C. G., VonDran, M. W., Stillman, A. A., Huang, Y., Hempstead, B. L., & Dreyfus, C. F. (2014). Astrocyte-derived BDNF supports myelin protein synthesis after cuprizone-induced demyelination. Journal of Neuroscience, 34(24), 8186–8196.

Geffin, R., Martinez, R., de las Pozas, A., Issac, B., & McCarthy, M. (2017). Fingolimod induces neuronal-specific gene expression with potential neuroprotective outcomes in maturing neuronal progenitor cells exposed to HIV. Journal of neurovirology, 23(6), 808–824.

Gusterson, R., Brar, B., Faulkes, D., Giordano, A., Chrivia, J., & Latchman, D. (2002). The transcriptional co-activators CBP and p300 are activated via phenylephrine through the p42/p44 MAPK cascade. Journal of Biological Chemistry, 277(4), 2517–2524.

Han, C., Shen, J. K., Hornicek, F. J., Kan, Q., & Duan, Z. (2017). Regulation of microRNA-1 (miR-1) expression in human cancer. Biochimica et Biophysica Acta (BBA)-Gene Regulatory Mechanisms, 1860(2), 227–232.

Kappos, L., Antel, J., Comi, G., Montalban, X., O’Connor, P., Polman, C. H., et al. (2006). Oral fingolimod (FTY720) for relapsing multiple sclerosis. New England Journal of Medicine, 355(11), 1124–1140.

Kerschensteiner, M., Gallmeier, E., Behrens, L., Leal, V. V., Misgeld, T., Klinkert, W. E., et al. (1999). Activated human T cells, B cells, and monocytes produce brain-derived neurotrophic factor in vitro and in inflammatory brain lesions: a neuroprotective role of inflammation? Journal of Experimental Medicine, 189(5), 865–870.

Khani-Habibabadi, F., Askari, S., Zahiri, J., Javan, M., & Behmanesh, M. (2019). Novel BDNF-regulatory microRNAs in neurodegenerative disorders pathogenesis: An in silico study. Computational biology and chemistry, 83, 107153.

Lee, D.-H., Seubert, S., Huhn, K., Brecht, L., Rötger, C., Waschbisch, A., et al. (2017). Fingolimod effects in neuroinflammation: Regulation of astroglial glutamate transporters? PloS one, 12(3), e0171552.

Li, E.-Y., Zhao, P.-J., Jian, J., Yin, B.-Q., Sun, Z.-Y., Xu, C.-X., et al. (2019). Vitamin B1 and B12 mitigates neuron apoptosis in cerebral palsy by augmenting BDNF expression through MALAT1/miR-1 axis. Cell Cycle, 18(21), 2849–2859.

Li, J.-H., Liu, S., Zhou, H., Qu, L.-H., & Yang, J.-H. (2014). starBase v2. 0: decoding miRNA-ceRNA, miRNA-ncRNA and protein–RNA interaction networks from large-scale CLIP-Seq data. Nucleic acids research, 42(D1), D92–D97.

Li, Q., Feng, Y., Chao, X., Shi, S., Liang, M., Qiao, Y., et al. (2018). HOTAIR contributes to cell proliferation and metastasis of cervical cancer via targetting miR-23b/MAPK1 axis. Bioscience reports, 38(1), BSR20171563.

Livak, K. J., & Schmittgen, T. D. (2001). Analysis of relative gene expression data using realtime quantitative PCR and the 2(-Delta Delta C(T)) Method. Methods, 25(4), 402–408, doi:10.1006/meth.2001.1262.

Luan, W., Li, L., Shi, Y., Bu, X., Xia, Y., Wang, J., et al. (2016). Long non-coding RNA MALAT1 acts as a competing endogenous RNA to promote malignant melanoma growth and metastasis by sponging miR-22. Oncotarget, 7(39), 63901.

Mazdeh, M., Monhaser, S. K., Taheri, M., & Ghafouri-Fard, S. (2019). A non-randomized clinical trial to evaluate the effect of fingolimod on expanded disability status scale score and number of relapses in relapsing-remitting multiple sclerosis patients. Clinical and translational medicine, 8(1), 11.

Mazibrada, G., Sharples, C., & Perfect, I. (2018). Real-world experience of fingolimod in patients with multiple sclerosis (MS Fine): An observational study in the UK. Multiple Sclerosis Journal–Experimental, Translational and Clinical, 4(4), 2055217318801638.

McTigue, D. M., Horner, P. J., Stokes, B. T., & Gage, F. H. (1998). Neurotrophin-3 and brain-derived neurotrophic factor induce oligodendrocyte proliferation and myelination of regenerating axons in the contused adult rat spinal cord. Journal of Neuroscience, 18(14), 5354–5365.

Modarresi, F., Faghihi, M. A., Lopez-Toledano, M. A., Fatemi, R. P., Magistri, M., Brothers, S. P., et al. (2012). Inhibition of natural antisense transcripts in vivo results in gene-specific transcriptional upregulation. Nature biotechnology, 30(5), 453.

Pahlevan Kakhki, M., Nikravesh, A., Shirvani Farsani, Z., Sahraian, M. A., & Behmanesh, M. (2018). HOTAIR but not ANRIL long non-coding RNA contributes to the pathogenesis of multiple sclerosis. Immunology, 153(4), 479–487.

Pelletier, D., & Hafler, D. A. (2012). Fingolimod for multiple sclerosis. New England Journal of Medicine, 366(4), 339–347.

Qi, K., & Zhong, J. (2018). LncRNA HOTAIR improves diabetic cardiomyopathy by increasing viability of cardiomyocytes through activation of the PI3K/Akt pathway. Experimental and therapeutic medicine, 16(6), 4817–4823.

Salmena, L., Poliseno, L., Tay, Y., Kats, L., & Pandolfi, P. P. (2011). A ceRNA hypothesis: the Rosetta Stone of a hidden RNA language? Cell, 146(3), 353–358.

Shi, B., Wang, Y., & Yin, F. (2017). MALAT1/miR-124/Capn4 axis regulates proliferation, invasion and EMT in nasopharyngeal carcinoma cells. Cancer biology & therapy, 18(10), 792–800.

Stadelmann, C., Kerschensteiner, M., Misgeld, T., BruÉck, W., Hohlfeld, R., & Lassmann, H. (2002). BDNF and gp145trkB in multiple sclerosis brain lesions: neuroprotective interactions between immune and neuronal cells? Brain, 125(1), 75–85.

Thompson, A. J., Banwell, B. L., Barkhof, F., Carroll, W. M., Coetzee, T., Comi, G., et al. (2018). Diagnosis of multiple sclerosis: 2017 revisions of the McDonald criteria. The Lancet Neurology, 17(2), 162–173.

Tian, N., Cao, Z., & Zhang, Y. (2014). MiR-206 decreases brain-derived neurotrophic factor levels in a transgenic mouse model of Alzheimer’s disease. Neuroscience bulletin, 30(2), 191–197.

Varendi, K., Kumar, A., Härma, M.-A., & Andressoo, J.-O. (2014). miR-1, miR-10b, miR-155, and miR-191 are novel regulators of BDNF. Cellular and molecular life sciences, 71(22), 4443–4456.

Wu, P., Zuo, X., Deng, H., Liu, X., Liu, L., & Ji, A. (2013). Roles of long noncoding RNAs in brain development, functional diversification and neurodegenerative diseases. Brain research bulletin, 97, 69–80.

Xiao, J., Hughes, R. A., Lim, J. Y., Wong, A. W., Ivanusic, J. J., Ferner, A. H., et al. (2013). A small peptide mimetic of brain-derived neurotrophic factor promotes peripheral myelination. Journal of neurochemistry, 125(3), 386–398.

Xu, L., Zhang, Z., Xie, T., Zhang, X., & Dai, T. (2016). Inhibition of BDNF-AS provides neuroprotection for retinal ganglion cells against ischemic injury. PloS one, 11(12), e0164941.

Yang, X., Wu, Y., Zhang, B., & Ni, B. (2018). Noncoding RNAs in multiple sclerosis. Clinical epigenetics, 10(1), 1–12.

Yao, J., Wang, X. Q., Li, Y. J., Shan, K., Yang, H., Yao, M. D., et al. (2016). Long non-coding RNA MALAT1 regulates retinal neurodegeneration through CREB signaling. EMBO molecular medicine, 8(4), 346–362.

Yuan, P., Cao, W., Zang, Q., Li, G., Guo, X., & Fan, J. (2016). The HIF-2α-MALAT1-miR-216b axis regulates multi-drug resistance of hepatocellular carcinoma cells via modulating autophagy. Biochemical and biophysical research communications, 478(3), 1067–1073.

Zhang, W., Qin, L., Wang, J., Fan, J., Lei, C., Liu, Q., et al. (2018). HOTAIR promotes proliferation, migration and invasion of esophageal squamous cell carcinoma by regulating MAPK1. INTERNATIONAL JOURNAL OF CLINICAL AND EXPERIMENTAL MEDICINE, 11(5), 4500–4511.

Zhou, K.-R., Liu, S., Sun, W.-J., Zheng, L.-L., Zhou, H., Yang, J.-H., et al. (2016). ChIPBase v2. 0: decoding transcriptional regulatory networks of non-coding RNAs and protein-coding genes from ChIP-seq data. Nucleic acids research, gkw965.

Zhou, Q., Chen, F., Zhao, J., Li, B., Liang, Y., Pan, W., et al. (2016). Long non-coding RNA PVT1 promotes osteosarcoma development by acting as a molecular sponge to regulate miR-195. Oncotarget, 7(50), 82620.

